# Hymenoptera (Insecta) telomerase RNAs switched to plant/ciliate-like biogenesis

**DOI:** 10.1101/2022.10.19.512496

**Authors:** Petr Fajkus, Matej Adamik, Andrew D.L. Nelson, Agata M. Kilar, Michal Franek, Michal Bubenik, Radmila Frydrychova Capkova, Alena Votavova, Eva Sykorova, Jiri Fajkus, Vratislav Peska

## Abstract

In contrast to the catalytic subunit of telomerase, its RNA subunit (TR) is highly divergent in size, sequence and biogenesis pathways across eukaryotes. Current views on TR evolution assume a common origin of TRs transcribed with RNA polymerase II in Opisthokonta (the supergroup including Animalia and Fungi) and Trypanosomida on one hand, and TRs transcribed with RNA polymerase III under the control of type 3 promoter, found in TSAR and Archaeplastida supergroups (including e.g., ciliates and Viridiplantae taxa, respectively). Here we focus on unknown TRs in one of the largest Animalia order - Hymenoptera (Arthropoda) with more than 300 available representative genomes. Using a combination of bioinformatic and experimental approaches, we identify their TRs. In contrast to the presumed type of TRs (H/ACA box snoRNAs transcribed with RNA Polymerase II) corresponding to their phylogenetic position, we find here short TRs of the snRNA type, likely transcribed with RNA polymerase III under the control of the type 3 promoter. The newly described insect TRs thus question the hitherto assumed monophyletic origin of TRs across Animalia and point to an evolutionary switch in TR type and biogenesis that was associated with the divergence of Arthropods.

**Graphical abstract:** 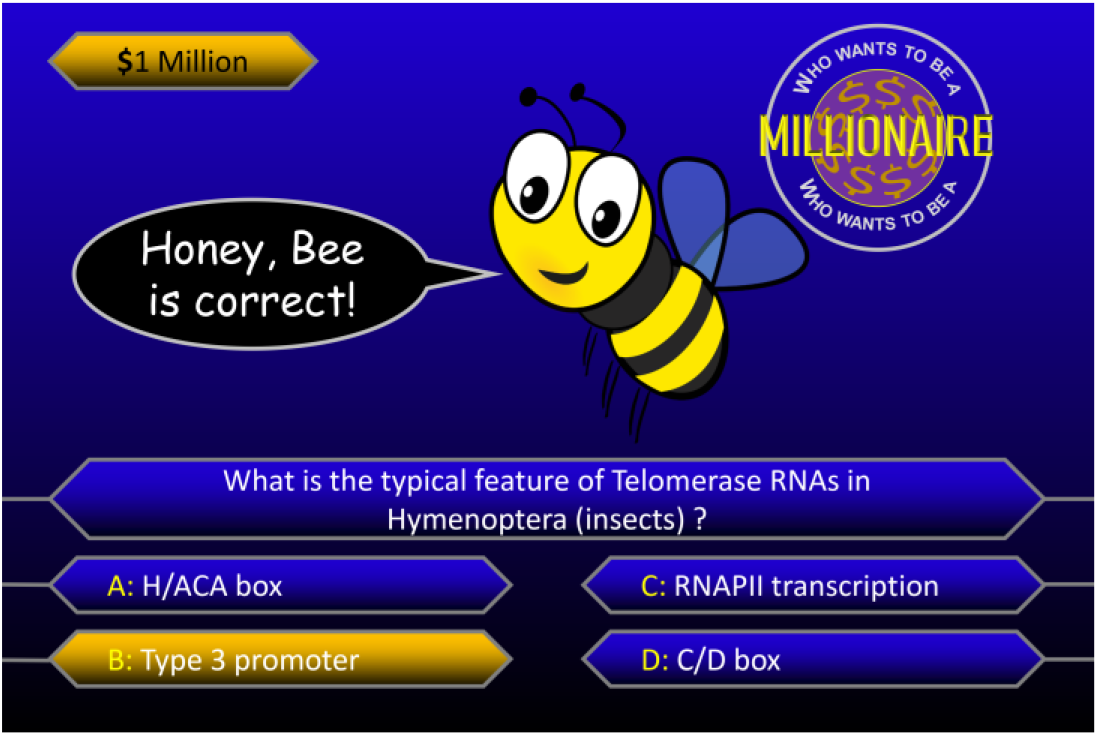

## INTRODUCTION

Telomerase, telomere DNA and associated proteins, which have evolved in concert, represent a well-tuned machine guarding genome stability in eukaryotic organisms, as solutions to the end-protection and end-replication problems. A change in any of its components can have serious consequences for cell and organism viability. Interestingly, specific mechanisms by which this machinery works in organisms from distant taxonomic groups are very diverse, including changes in the telomere DNA sequence, telomere accessory proteins, distinct telomerase biogenesis and regulation pathways. In rare cases, a telomerase-independent telomere maintenance system was established (reviewed in (1)). A part of this diversity stems from the most enigmatic telomerase component – the Telomerase RNA (TR). TR serves as a scaffold for the entire telomerase ribonucleoprotein complex assembly, but most importantly, TR provides a short template region for synthesis of telomere DNA repeats at the ends of chromosomes. This is achieved using the catalytic protein component Telomerase Reverse Transcriptase (TERT). TRs are extremely divergent in size, sequence, structures and types of non-coding RNA (ncRNA) biogenesis pathways (reviewed in (2)), which have made the identification and characterization of TRs extremely difficult. Thanks to the expanding availability of sequenced genomes, TR genes were computationally predicted and experimentally characterized in plants (3), green algae and some members of the TSAR supergroup (4), and yeasts (5) as well as early diverged Animalia species. These include Protostomia, Placozoa and Porifera (6). Current views on TR evolution proposed by us and others (4,6,7) assume a common origin of TSAR and Archaeplastida TRs (as RNAPIII transcripts) on one hand, and Opisthokonta (Fungi + Animalia) and Trypanosomida (as RNAPII transcripts) on the other (**Figure 1**). Previously, the first characterized TRs in Ciliates (TSAR) (8), as short snRNAs with the type 3 snRNA RNAPIII promoter (9,10), were considered as evolutionary outliers compared to the considerably longer RNAPII-transcribed TRs in animals and fungi. However, newly characterized TSAR and Archaeplastida TRs, with the same snRNA promoter as ciliate TRs, indicate that RNAPIII transcription of TRs is probably ancient, dating back to the divergence of TSAR and Archaeplastida (4). Considering the currently recognized topology of the root of eukaryotic phylogeny (11), and recently acquired knowledge of TRs, there is no unambiguous answer to the question of whether the first TR originated as an RNAPII or RNAPIII transcript. (**Figure 1**). The proposed scenario of the monophyletic origin of Archaeplastida + TSAR TRs as type 3 promoter RNAPIII transcripts, vs. Animalia TRs as H/ACA box RNAPII transcribed small nucleolar RNAs (snoRNAs), is well supported by comprehensive sets of newly characterized TRs from evolutionarily distant and early divergent organisms, which demonstrated conservation of typical TR secondary structural elements ((4,6)). In other words, both studies (4,6) inferred novel TR genes in several groups from Archaeplastida + TSAR or Animalia, based on some previous knowledge of small nuclear RNA biogenesis and regulation (e.g. type 3 RNAPIII promoters in plants and ciliates, or H/ACA snoRNAs in Vertebrates + Echinoderms). However, TR inference failed in both studies for many large and deeply sequenced taxonomic groups (e.g., Oomycota, Rhodophyta, Apicomplexa or Arthropods and Nematodes) inside the aforementioned eukaryotic kingdoms. Several reasons why computational strategies had failed were proposed, e.g., enormous TR variability beyond the computational limits to infer their evolutionarily conserved sequence motifs or structural features. Another possibility could be transitions in characteristic features of TRs and their genes, which might have happened in association with the divergence of some new taxa. These evolutionary transitions would then result in switches between TR promoters, transcription factors, RNAPII and RNAPIII transcription, and structural features of TRs (**Figure 1**). In this study, we focus on unknown TRs in one of the largest Animalia orders - Hymenoptera (Arthropoda) with more than 300 available representative genomes. Based on our data presented in this study, published work and preprint ((12,13)), hymenopteran species show remarkable telomere DNA sequence variability. As the most plausible scenario of telomere sequence changes in Hymenoptera, we assume corresponding changes in TR template regions dictating the sequence synthesized by telomerase. Thus, the mechanistic explanation of telomere DNA evolutionary changes in Hymenoptera can be provided by TR identification across their phylogeny. Here, we decipher insect TRs *de novo* and unexpected features of Hymenoptera TRs that have not been observed in Animalia so far. These TRs are short snRNAs, and likely transcribed by RNAPIII under the control of the type 3 promoter. Such features are characteristic for TRs in Archaeplastida + TSAR and challenge the current concept of monophyletic origin of TRs in animals.

**Figure 1.**
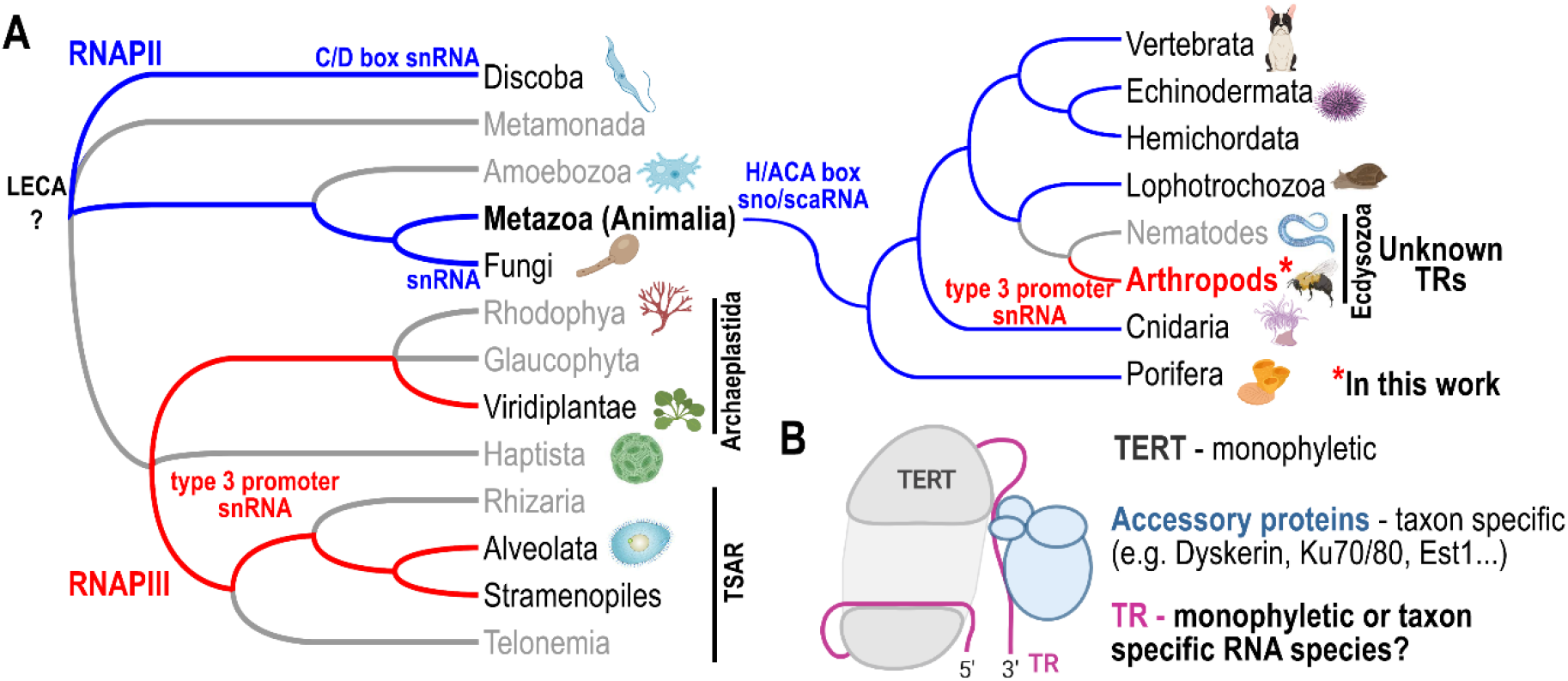
Current view on divergent TR evolution. (**A**) Different TR biogenesis pathways exclusively employed in separate evolutionary lineages are highlighted. TRs transcribed with RNAPII (C/D box snRNAs in trypanosomes; H/ACA box snoRNAs in Animalia, snRNA in yeasts) are depicted in blue. TRs generated as RNAPIII-transcribed snRNAs with the type 3 promoter in Archeplastida + TSAR are highlighted in red. An unexpected discovery of Hymenoptera TRs (Arthropoda) as the type 3 promoter snRNAs, presumably transcribed by RNAPIII, radically changes the previous view on monophyletic TR evolution in Animalia. (**B**) Simplified illustration of the telomerase complex as an interplay of at least three components – a conserved (monophyletic) protein subunit (TERT), telomerase RNA subunit (TR) the subject of this study, and taxa-specific accessory proteins. The common origin of animal TRs is questioned by insect TRs identified in this study.

## MATERIALS AND METHODS

### Identification of telomere motifs in Hymenoptera

The screening for telomere motifs in *Hymenoptera* was done in species with publicly available genome assemblies in NCBI. We used Tandem Repeats Finder – TRFi (14), and custom-made scripts (15) to analyse raw sequencing data downloaded from the Sequence Read Archive (SRA) at NCBI. Paired-end datasets from Illumina platforms with at least 2.5 million spots per species were preferred for analysis (i.e., 5 million paired-end reads per species were typically analysed). The output of TRFi analysis was sorted (in descending order) and taxonomically categorized in families or superfamilies. To predict candidate telomere sequences, the most frequent tandem motifs were checked for: i) the corresponding region in TR template region; ii) common motifs with relatives; iii) knowledge of telomere sequences from literature or chromosome/scaffold ends in respective genome assemblies. In examples from Cynipoidea, candidate telomere sequences were predicted using different approaches. In the chromosome level genome assembly of *Belonocnema kinseyi* (GCA_010883055), 2kb-long terminal sequences were used in reciprocal blastn searches to uncover common sequences among chromosome ends. In *Leptopilina boulardi* we predicted telomere candidates as the most frequent repeats at the termini of genome assembly (GCA_019393585) scaffolds. *L. boulardi* repeats were identified *de novo* in RepeatExplorer2/TAREAN (16) upon a set of short Illumina pair-end genomic reads (SRA accession: SRR11665922, parameters: tblastx; Word size=3; Cluster size threshold=0.001, and Minimal cluster size for assembly=5). Using RepeatExplorer2 “Assembly Annotation Tools” with default settings, repeats were annotated in the genome assembly according to similarity with *de novo* identified Clusters. Terminal sequences (1 kb each, repeat annotation included) from all reference genome contigs were extracted. The extracted annotations were sorted according to their frequency in the terminal sequences. For the top 20 most frequent clusters, detailed data are presented (e.g., consensus sequence, dotplots, annotation) (Supplementary Table S5), full RepeatExplorer2 output data are available upon request.

### TR prediction in Bombus genomes

Candidate TR loci were initially predicted in Geneious Prime 2020.2.5 (https://www.geneious.com) in 12 available chromosome-level *Bombus* representative genome assemblies at NCBI (listed in **Supplementary table S1)** as 400 nt genomic sequences with a template-like region in the middle. As the template-like region, we considered both strands of any circular permutation of the corresponding telomere repeat (i.e. TTAGGTTGGGC in *B. sylvestris*, and TTAGGTTGGGG in other 11 *Bombus* genomes) + 1 nt as a minimal annealing portion of the template. Sites with tandem arrays of three and more motifs (telomeric or interstitial telomere repeats) were excluded from the TR-like loci search. Overlapping TR-like loci were considered as a single candidate locus. Follow up TR loci datasets per genome were utilized in mutual blastn searches (default scoring/parameters; threshold ≤ 1e-20) to identify shared TR candidates among Bombus genomes. Shared TR candidate/s were further utilized in blastn searches in other 18 scaffold/contig level *Bombus* genome assemblies (incl. *B. waltoni* and *B. superbus* with TTAGG resp. TTTAGGG telomere repeats) and other genomes outside the *Bombus* genus (with threshold ≤ 1e-5).

### TR homology searches in Hymenoptera

Predicted transcribed regions of *Bombus* TR candidates and related homologs from close relatives identified by blastn (e.g. in *Tetragonula sp*.) were utilized for construction of the first covariance model (CM) by Infernal 1.1.2 tool (17) for sequence-structure homology searches of TR homologs. TR transcription boundaries and transcript orientation were estimated in *B. terrestris* according to the mapping of total RNAseq data (SRR5614374) to the *B. terrestris* genome using Read Mapping and Transcript Assembly workflow (RMTA) (18). TR sequences were aligned and folded using the LocARNA package (19) as an input for the construction of the CM by the Infernal tool. Novel hits produced by Infernal searches in Hymenoptera representative genomes were checked according to their significance (e-value), the presence of corresponding template regions and promoter structures (Infernal hits were extended with 200 nts of genomic context). CMs were progressively optimised with newly “verified” TR hits from repeated Infernal searches. To increase the efficiency of TR Infernal searches across divergent Hymenoptera species, CMs were optimised according to phylogeny relationships for specific taxons (e.g. Apoidea, Formicoidea, Chalcidoidea and others)

### Identification of type 3 snRNA promoters in Arthropoda

Type 3 snRNA promoters were characterised similarly as in (4) with the following adjustments. The promoters were sought in all available representative genome assemblies from Arthropoda (without Diptera) in NCBI using Infernal with CMs available in the RFAM database (20). These included CMs for U1 (RFAM no.: RF00003); U2 (RF00004); U3 (RF00012); U4 (RF00015); U5 (RF00020); U6 (RF00026): U6atac (RF00619); MRP (RF00030); 7SK (RF01052) RNAs. Output sequences were extended for 200nt of genomic context to obtain their putative “promoter” regions and filtered according to hit significance (e-value). Promoter regions from significant hits were screened for the presence and topology of common sequence motifs using the MEME tool (21) or manually by their multiple sequence alignments within particular organisms. Results from type 3 promoter screenings in Hymenoptera and Lepidoptera genomes were compared with previously characterized type 3 promoters in *Apis mellifera* (Hymenoptera) and *Bombyx mori* (Lepidoptera) (22).

### Search for TERT across Hymenoptera genomes

Full-length TERT amino acid sequences from 22 species across the Hymenoptera phylogeny (shown in **Supplementary Figure S2**) were collected from InsectBase 2.0 (23). These sequences were split into two parts containing the TERT-specific Telomerase RNA Binding Domain (TRBD) or the Reverse Transcriptase domain (RT) that is related to retroviral reverse-transcriptases. TERT sequences were used as queries in tblastn searches (Word size = 3; Gap costs = 11/1; E-value threshold = 0.00001) for TERT-like DNA sequences in Hymenoptera representative genomes at NCBI. Tblastn results were sorted according to respective genomes with the Tidyverse package (www.tidyverse.org), and visualised as a heatmap according to e-value (resp. its logarithmic scale) using ggplot2 (24).

### Search for TR-like genes with type 3 promoter in Lepidoptera

TR candidates in Lepidoptera genomes were predicted similarly as described in (4). Specifically, all type 3 promoter–like sequences (obtained from above, “*Identification of type 3 snRNA promoters in Arthropoda*”) were annotated in example genomes (*Spodoptera exigua* and *Plutella xylostella*) with highly conserved promoter motif elements. Promoter-like sequences were extended with 400nt long genomic context downstream and checked for a presence of the template-like region (i.e., circular permutation of a cytosine rich strand of the telomere motif + one nucleotide as a minimal annealing portion (i.e. AACCTA/ACCTAAC/…/TAACCT). TR candidates were used in homology searches (blastn) in other Lepidoptera representative genomes. As relevant TR-like homologs in blastn searches, we considered only sequences that showed: i) TR sequence similarity; ii) type 3 promoter; iii) template region corresponding to the telomere repeat.

### TR structure prediction

TR structures were inferred from multiple sequence alignments (MSAs) generated from all putative TR sequences in each major lineage (Meliponini, Bombini, Halictidae, Vespoidea, Pteromalidae, Tenthredinoidea, and Lepidoptera; see **Supplementary Figure S4** for specific taxa and sequences). MSAs were generated in Geneious (v11.0.14.1; https://www.geneious.com) using MAFFT with default parameters. To identify putative pseudoknot elements, long range interaction regions surrounding the template were identified within each MSA separately using conservation or covariation as a guide (demarcated as the core-delimiting P1C). This delimited region surrounding the template was examined for pseudoknot elements with a particular focus on anchor points (conserved elements within the region) within the MSA.

### TRAP assay

Up to 50mg of gonadal tissue (ovaries) were extracted using of 200 µl of extraction buffer accordinq to (25), (10 mM Tris/HCl pH7,5, 1 mM EGTA, 0,1 mM benzamidine, 5 mM 2-mercaptoethanol, 0,5 % (w/v) Chaps, 10 % (v/v) glycerol, 40 U/ml RNasin® Ribonuclease Inhibitor (Promega)). The tissue was homogenized by pestle in a 1.5 ml Eppendorf tube and then incubated on ice for 30 min. The debris was removed by centrifugation (20000g, 20min, 4 °C) and the supernatant was aliquoted (20 µl) and stored at −80 °C. The protein concentration of extracts was measured by Pierce™ Coomassie Plus (Bradford) Assay Kit (Thermo Fisher Scientific) and amounts corresponding to 0.5 µg and 5 µg of protein were used as an input to the TRAP assay reactions, in which 5 pmol of substrate primer and 12,5 µl of TB Green Premix Ex Taq II (Takara) were added. First, the telomerase reaction was performed at 30 °C for 60 min, then 5 pmol of the reverse primer was added and 30 cycles of the PCR was performed (94 °C 30s, 60 °C 30s), with the final extension at 72 °C for 10min). The PCR products were separated on 10% polyacrylamide, 0.5x TBE gels in a Mini-PROTEAN Tetra Vertical Electrophoresis Cell using 100V, 85-90min. The gels were stained with SYBR™ Gold (Invitrogen) and visualized on a Typhoon FLA 9000 imager (GE Healthcare).

### Chromosomal preparations and FISH

Chromosomal spreads were made from gonads of adult individuals from species listed in **Supplementary Table S6** according to (26). Microscopic slides were rehydrated in 2x SSC, then incubated in 10 ug/ml RNAse A for 30 minutes at 37°C in a humid chamber. After washing in 2x SSC, slides were dehydrated in an ethanol series (70% - 80% - 90% ethanol, 2 min each). Slides were then air-dried and hybridization mix (10% dextran sulfate, 50% deionized formamide in 2x SSC + 10 nM custom telomere LNA probes (see **Supplementary Table S6**)) was pipetted onto the slides. Slides were denatured on a hot plate (80°C, 4 min), and incubated in a humid chamber overnight at 37°C. The next day, slides were washed in 2x SSC for 5 min, then 3 times in 50% formamide in 2x SSC and 42°C, then dehydrated in an ethanol series (same as above) and mounted in DAPI (Invitrogen) + VECTASHIELD (Vector laboratories). Samples were imaged on a Zeiss AxioImager Z2 microscope with 63x (1.4 NA) Plan Apochromat oil objective, using DAPI, TRITC and Cy5 filters, illuminated with a UV-lamp to detect DNA staining by DAPI and probes with Cy3 and Cy5 fluorescent tags. Images were captured on a Hamamatsu ORCA Flash camera with a 2048 × 2048 pixel resolution.

### Northern hybridizaton

Total RNAs were isolated by TRI Reagent® (Molecular Research Center) from approx. 50mg of female gonads of *B. terrestris* and *A. mellifera*. RNA concentrations were measured by NanoDrop™. Northern hybridisation experiments were carried out as described in (4) with [32P-dATP]-labelled PCR probes from *B. terrestris* and *A. mellifera* TRs. AmTR and BtTR were amplified from genomic DNA of *A. mellifera* and *B. terrestris* and cloned by TOPO™ TA Cloning™ Kit (Invitrogen). PCR primers were as referred to in **Supplementary table S6**.

## RESULTS AND DISCUSSION

The only known or easily predictable feature of a completely unknown TR is its template region, which has to correspond to the sequence of the telomere repeat. In this work we utilised a similar strategy as in the case of the unusually long telomere repeat in *Allium* sp. (CTCGGTTATGGG) (27), which allowed us to predict the first plant TRs in transcriptomic data and subsequent identification of the TR homologs across the whole plant phylogeny (3). Here we predicted and utilized long and variable telomere repeats in bumblebees (*Bombus* sp.) in order to identify putative TR genes in the genome assemblies and infer their TR homologs towards evolutionarily distant species. Compared to the previous studies inferring TRs in early diverged Animalia or Archaeplastida + TSAR, based on assumptions formulated on the basis of already known TRs (4,6), here we decipher insect TRs *de novo*, equipped only with a knowledge of telomere sequences. The comprehensive set of Hymenoptera TRs and corresponding telomere repeats presented in this study are not just a mere “collecting” of additional Animalia TRs or repeat motifs, although it may seem so. Hymenoptera TRs showed unexpected features, such as short snRNAs plausibly transcribed by RNAPIII under the control of the type 3 promoter. This is also characteristic of TRs in Archaeplastida + TSAR, but has not been observed in Animalia so far. Thus, instead of results in accordance with the monophyletic concept of Animalia TRs as H/ACA box snoRNAs transcribed by RNAPII (6), Hymenoptera TRs point to surprising evolutionary switch in TR biogenesis.

### Telomere motif variability across Hymenoptera

Insecta exhibit unprecedented diversity in telomere/telomerase evolutionary pathways, with both canonical telomerase-based mechanisms of telomere maintenance and various telomerase-independent mechanisms that evolved after the loss of telomerase in Diptera (1,28)). Moreover, evolutionary switches in telomere DNA sequences were recently identified in Coleoptera (29) and Hymenoptera (12,13). Without insects, the evolution and diversity of telomere repeats and associated TR template regions in animals would be monotonous compared to, e.g., those in yeasts or plants (30,31). Here we have expanded and confirmed recently predicted (12,13) telomere motifs in Hymenoptera species (**Figure 2, Supplementary Table S1 and S2**). This was achieved by using a modified Tandem Repeats Finder tool (TRFi) (14) for identification of telomere repeats in raw genomic data, which we have utilised for this purpose several times in various eukaryotic organisms (27,29,32,33). TRFi results on raw genomic data are limited to species with available genomes and summarized in **Supplementary Table S2** (note: The first sheet contains hyperlinks to particular TRFi results sorted according to species taxonomy). Besides TRFi, we exploited the availability of numerous high-quality telomere-to-telomere genome assemblies to verify predicted telomere motifs at chromosome/scaffold ends, similarly as was utilized in (12) (**Figure 2**).

**Figure 2.**
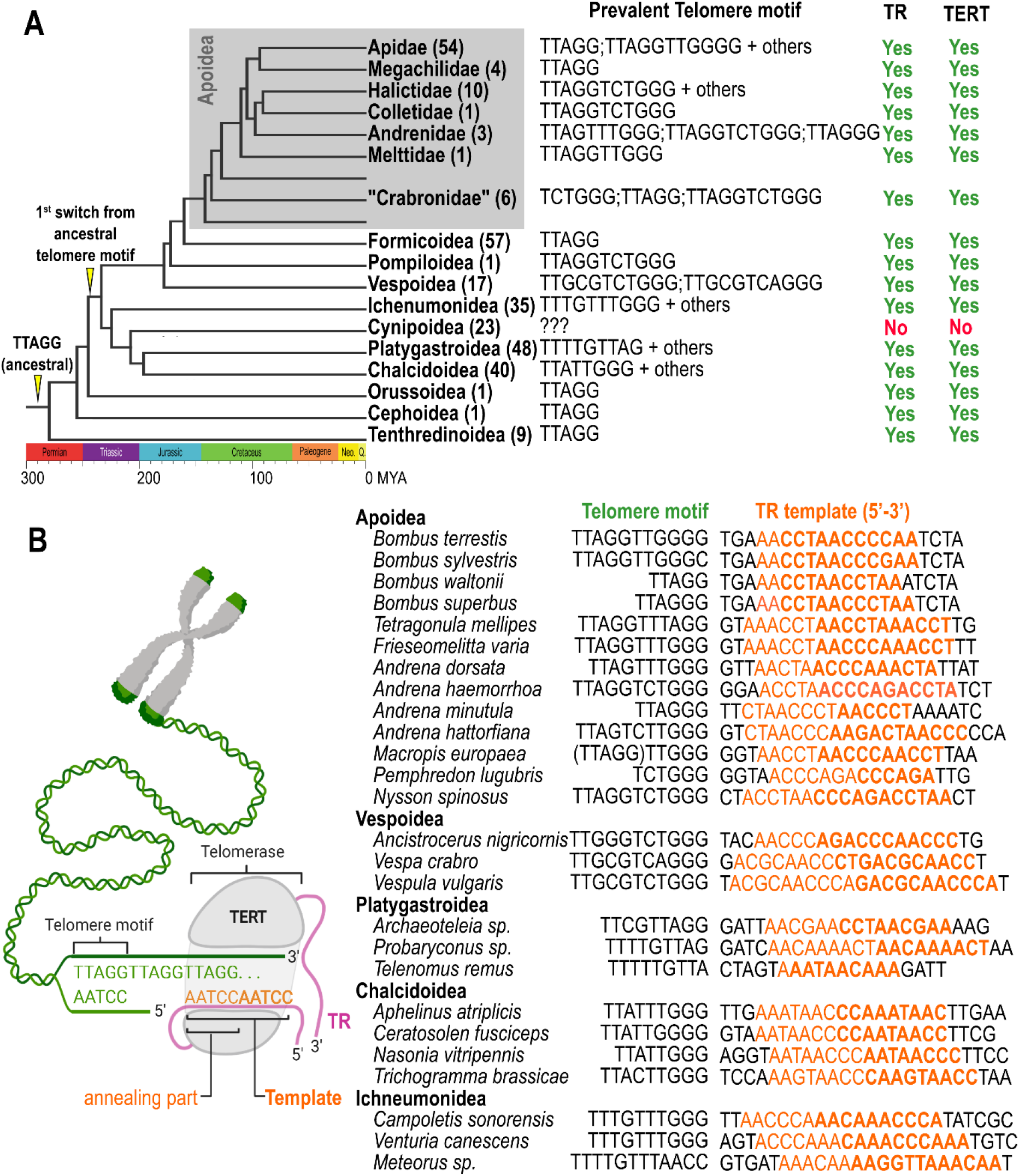
Telomere motifs and telomerase RNAs in Hymenoptera. (**A**) Simplified overview of newly characterized and previously predicted (12,13) telomere motifs in Hymenptera phylogeny. Several available representative genomes are indicated for respective clades. A topology of presumed evolutionary switch from the ancestral TTAGG repeat in insects is marked on the phylogeny tree. Taxa in which we predicted TRs and TERT sequences are marked in TR/TERT columns. Analysis of 23 Cynipoidea genomes did not reveal candidate TR/TERT genes and various sequences were identified in terminal regions of assembled genomes (details in text, **Supplementary Figure S3** and **Supplementary Table S5)**. (**B**) Examples showing independently predicted TRs (resp. their template regions) and telomere motifs (*in *Bombus* sp., telomere motifs were employed for initial TR sequence prediction). Mutually corresponding changes in telomere motifs and template regions of TR homologs (the right part), especially among closely related species (e.g. *Andrena* sp.) greatly support the validity of presented TRs/telomere motifs, as schematically depicted on the left. Comprehensive data for respective species are shown in **Supplementary Tables S1** and **S2**.

In addition, the previous work studying telomeres and telomerase activity in *Bombus terrestris* reported an ancestral TTAGG telomere repeat (34). However, we experimentally confirmed an unusually long telomere repeat TTAGGTTGGGG predicted by us and (12) (**Figure 3**) using *in situ* hybridisation, cloning and sequencing of the products from the Telomere Repeat Amplification Protocol (TRAP). The discrepancy with (34) can be explained by the fact that TTAGG is a sub-repeat of the genuine telomere repeat in *B. Terrestris*, which can result in cross-hybridisation and false positive results in methods used in the previous work (i.e., quantitative real-time TRAP and Southern hybridisation). Correspondingly, the TTAGG motif was also irregularly concatenated in our sequenced clones of TRAP PCR products formed primarily by TTAGGTTGGGG arrays (**Supplementary Figure S1**).

**Figure 3.**
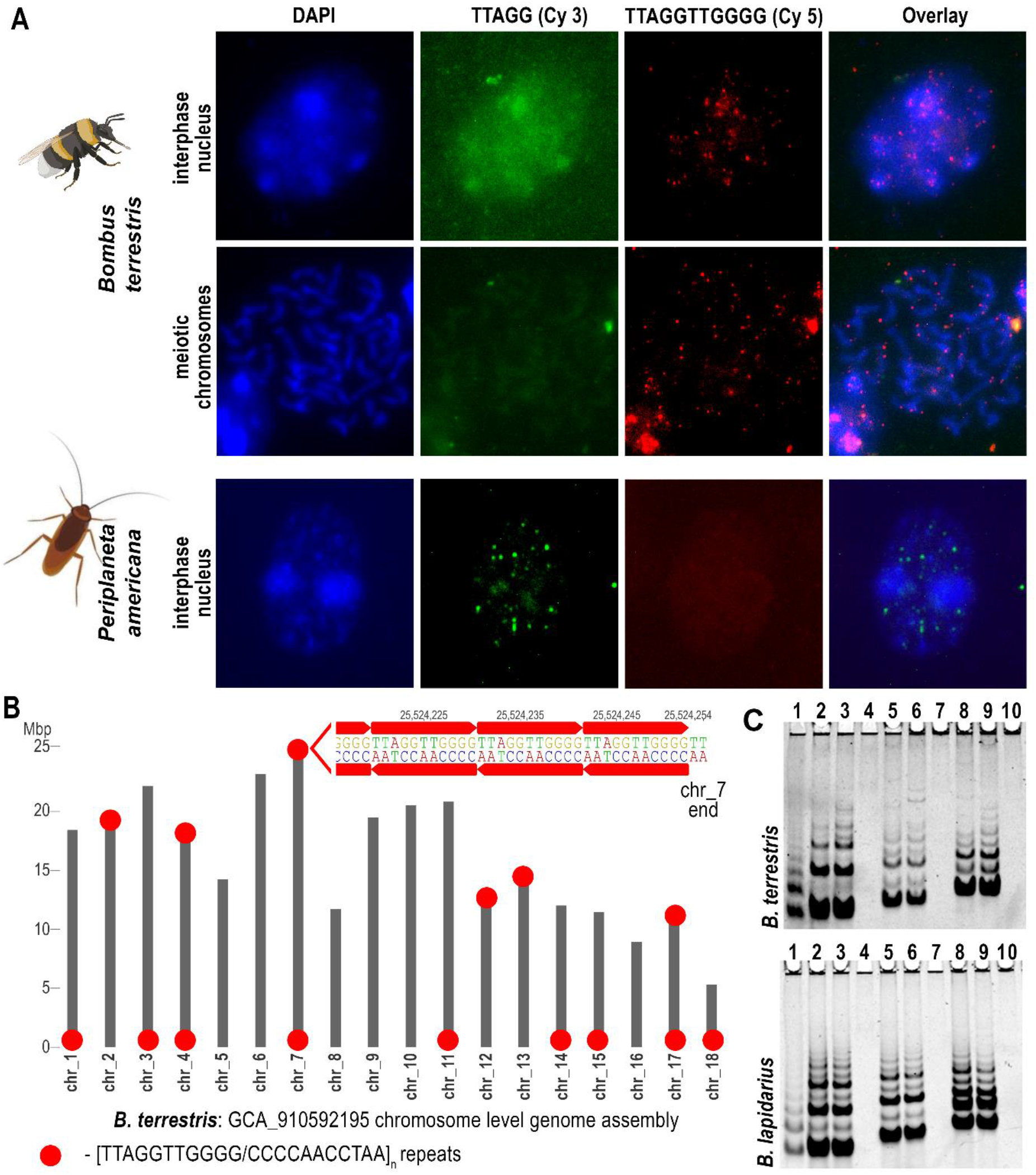
Validation of telomere sequence in *Bombus*. (**A**) Fluorescence *in situ* hybridization of telomere probes on interphase nuclei and meiotic chromosomes in *B. terrestris* and *P. americana*. The probe with the typical insect motif TTAGG (green) was directly labelled with Cy3 and the *Bombus* sequence TTAGGTTGGGG (red) was labelled with Cy5. The chromosomes and nuclei of *B. terrestris* provided a specific signal only with the TTAGGTTGGGG probe, while the control sample of *Periplaneta americana* hybridized specifically only with the TTAGG probe. (**B**) Analysis of terminal regions of chromosomes in *B. terrestris* genome assembly. Presence of TTAGGTTGGGG/CCCCAACCTAA tandem repeats at respective chromosome ends is depicted by red circles. (**C**) Detection of telomerase activity in *B. terrestris* (upper panel) and *B. lapidarius* (lower panel) using the TRAP assay and various combinations of substrate and reverse primers. Products were separated by PAGE and visualized with SYBR™ Gold. The oligonucleotides (substrate primer, reverse primer) and the volume of the protein extract are as follows: 1 – positive control *Periplaneta americana* (VRP440, BmCXa, 0.2ul), 2 - (VRP440, VRP437, 2ul) 3 - (VRP440, VRP437, 1ul) 4 - (VRP440, VRP437, negative control without telomerase extract) 5 - (TS21GGG, VRP437, 2ul), 6 - (TS21GGG, VRP437, 1ul), 7 - (TS21GGG, VRP437, negative control without telomerase extract), 8 - (47F, VRP437, 2ul), 9 - (47F, VRP437, 1ul), 10 - (47F, VRP437, negative control without telomerase extract). For primer sequences, see **Supplementary Table S6**.

Except for *Bombus* sp. (where the telomere motifs were utilised for prediction of TR candidates –see the paragraph “*TR prediction in Bombus and searches for TR homologs*”), all other predicted motifs were in perfect accordance with the template regions of independently inferred TRs, strongly supporting the correctness of predicted telomere repeats and TRs reported in this work (**Figure 2B, Supplementary Table S1 and S2**). Remarkably, both the TR and the telomere motif predictions failed in the deeply sequenced Superfamily Cynipoidea (23 assembled genomes). Moreover, chromosome level assembly from *Belonocnema kinseyi* or well assembled genomes from *Leptopilina boulardi* did not reveal any telomere-like repeats (short tandem arrays) at the chromosome/scaffold ends, as was found in most other Hymenoptera assemblies. Thus, we searched available Hymenoptera genomes for candidate *TERT* sequences to identify a putative protein subunit of telomerase that would suggest/support a telomerase-based system of telomere maintenance. The *TERT* loss in Cynipoidea would indicate a new case of telomerase-independent telomere maintenance, similar to what is known in Diptera evolution (28). When using tblastn with 22 full length TERT protein queries across the Hymenoptera phylogeny (**Figure 2B, Supplementary Figure S2A**), a TERT-specific TRBD domain and an RT domain distantly related to the other reverse transcriptases, TERT-like sequences were easily identifiable across all taxonomic groups (**Figure 2B**, in detail - **Supplementary Figure S2B**). TERT was missing/not detected in rare cases scattered through the Hymenoptera phylogeny (e.g., no hit in *Vespula pensylvanica* genome assembly vs. significant TERT hits in other closely related *Vespula* sp. genomes). This makes it inconclusive whether the TERT is really missing or the genome assembly is incomplete in these isolated cases. In contrast, the absence of TERT sequences was common to all 23 species from Cynipoidea (Figitidae + Cynipidae), suggesting an evolutionary switch to telomerase-independent telomere maintenance (**Figure 2A**, in detail **Supplementary Figure S2B**). In our effort to unveil telomeres in Cynipoidea, we inspected the chromosome level assembly of *Belonocnema kinseyi* (GCA_010883055, 10 chromosomes + 5510 unplaced scaffolds) and high quality genome assembly from *Leptopilina boulardi* (GCA_019393585, 409 scaffolds). In the case of *B. kinseyi* (no raw genomic data were available) we checked by blastn whether any particular chromosomes share any common terminal DNA. In *L. boulardi* raw genomic Illumina data (SRR11665922) were processed by RepeatExplorer2 (16) to identify and quantify repeats *de novo*. Repeats were annotated in the genome assembly to uncover their presence and frequency at various scaffold ends. We proposed a candidate telomere sequence in *B. kinseyi* as a 62 nt long satellite repeat present at the ends of chromosmes 1, 3, 8 and 10 (from total 10 chromosomes) (see **Supplementary Figure S3**). In *L. boulardi* we present the top 20 most frequent repeats (Clusters) identified *de novo* at the terminal positions (see **Supplementary table S5**). Specifically, 12 of 20 clusters represent various satellite repeats, two clusters correspond to simple repeats and Class II mobile elements, one to 45S rDNA and the Class I mobile element Helitron, and one repeat remained unresolved. With regard to *L. boulardi* chromosome number (n=10; i.e. 20 ends) and genome assembly generated from long reads (PacBio Sequel), which usually achieve optimal repeat resolution compared to assembly from short read sequencing, it is probable that telomere sequence(s) are present among our predicted candidates. However, predicted candidate telomere repeats as well as a plausible non-telomerase telomere maintenance system in Cynipoidea need to be further investigated experimentally.

### TR prediction in Bombus and searches for TR homologs

The search for telomere repeats in *Bombus* species comprised analysis of 30 well assembled and relatively small, closely related genomes (around 230 Mb). In addition to an unusually long telomere repeat, TTAGGTTGGGG, characterized in *B. terrestris* (see above), more telomere variants were found: TTAGGTTGGG**C** *in B. sylvestris*, TTAGG in *B. waltonii* and TTAGGG in *B. superbus*. Genomes harbouring long telomere repeats were utilized to identify relatively low numbers of putative TR loci in *Bombus* genomes. Except for *B. waltonii* and *B. superbus* (with much shorter repeats), *Bombus* genomes contained approx. 400 TR template-like loci per genome.

Template-like regions extended with ±200 nt of genomic context were used in reciprocal blastn searches among *Bombus* species to identify common TR candidate(s) that harbour a corresponding template. Thanks to the point mutation in the *B. sylvestris* telomere motif, (and thus in its corresponding template region), such filtering of candidate TRs was highly efficient. For example, from the total of 150 template-like regions in *B. sylvestris* (GCA_911622165), only 11 had a homolog among a total of 435 template-like regions in *B. terrestris* (GCF_000214255.1)). When the number of *Bombus* species was expanded with other chromosome level assemblies, only a **single** sequence common within the *Bombus* genus showed a TR template corresponding to the telomere repeat (see **Figure 2B** for examples). Follow up blastn homology searches against Hymenoptera genomes with the *Bombus* TR candidate as a query showed sequence homologs in closely related *Tetragonula* species. Their putative template regions corresponded to the predicted TTTAGGTTAGG telomere motif **(Figure 2B, Supplementary Table S1**), which supported the predicted candidate as the genuine Telomerase RNA. Transcribed regions of candidate TRs (see the paragraph *“TR transcription”*) were utilised for the construction of the first Covariance Model (CM) for the Infernal tool, which is much more robust for identification of structural RNA homologs compared to a standard sequence similarity search like blast. Reiterative Infernal searches with optimised CMs with novel TRs allowed us to gradually elucidate TR homologs across the Hymenoptera phylogeny (**Supplementary Table S1**).

### TR transcription

Since predicted Hymenoptera TRs are derived from genomic data, we verified their presence at the RNA level first in examples of *Apis mellifera* and *Bombus terrestris*. The first searches using available mRNA (poly(A) selected) data did not indicate clear transcription of the TR loci in either *A. mellifera* or *B. terrestris* (tested datasets are referred in **Supplementary Table S6**). However, available total RNA-seq data employing rRNA depletion showed a specific peak of coverage by RNA-seq reads, which corresponded to the lengths of RNAs detected by northern hybridisation with TR probes in the total RNA preparations in this work. (**Figure 4**).

**Figure 4.**
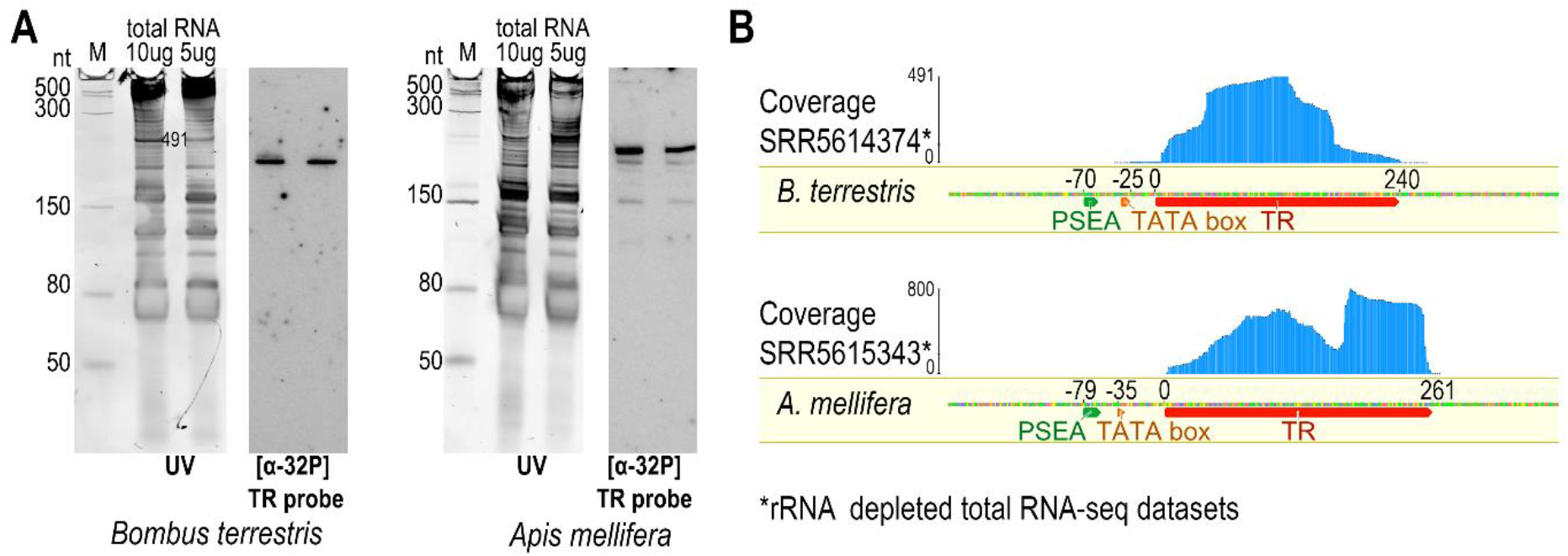
Verification of candidate TR transcripts in *B. terrestris* and *A. mellifera*. (**A**) Northern hybridization using corresponding radioactively labelled TR probe ([α-^32^P]dATP). Gels were stained with SYBR™ Gold and visualized in UV light (UV). Low Range ssRNA Ladder (NEB) was used as a marker (M). (**B**) - Mapping of total RNA-seq data (employing rRNA depletion) to the TR locus. RNA coverage indicates TR transcription and transcript boundaries. TR transcript and subsequently characterized type 3 promoter are indicated below the sequences.

Transcriptomic data allowed us to assess 5’ and 3’ transcript boundaries, as well as the correct transcript orientation and the promoter region. The rare occurrence of *Bombus* and *Apis* TRs in tested mRNA-seq datasets (**Supplementary Table S6**) may indicate the lack of poly(A) at the 3’ end, suggesting a switch from RNAPII transcription (that has been presumed for all Animalia TRs so far) to different RNA polymerases. This result is very similar to that obtained in plant and ciliate TRs that are transcribed with RNAPIII under the control of a type 3 promoter (3,4).

### Type 3 snRNA promoter defines genuine Hymenopteran TRs

Due to this parallel with plant and ciliate TRs, we compared Hymenopteran TR promoter regions with an already known type 3 snRNA promoter in *Apis mellifera* (22) and our newly predicted type 3 promoters for other species (**Supplementary Table S3**, examples in **Figure 5B**). In general, type 3 promoters are composed of a conserved species/taxon-specific sequence (usually referred to in literature as the Proximal/Upstream Sequence Element (PSE/USE)), and the TATA box. They can drive both RNAPII and RNAPIII transcription. RNAP specificity of the type 3 promoters is given by specific promoter features that vary among different taxa. For example, the specific distance between USE and TATA determines whether RNAPII or RNAPIII transcription starts at these loci in land plants, while TATA box strength/presence or absence is decisive in humans (reviewed in e.g. (35)). Type 3 promoters in insects were characterized as cis-acting elements with conserved PSEA and variable PSEB and TATA box. RNAPII promoters are composed of PSEA and PSEB, while RNAPIII promoters consist of only PSEA and a TATA box (22) (**Figure 5A**). Thus, a comparison of the candidate TR promoters with type 3 promoters of various snRNAs (**Supplementary Table S1 and S3**) confirmed the presence of a type 3 promoter in all TRs identified. Moreover, comparing examples across the Hymenoptera phylogeny **(Figure 5B)** shows that TR promoters are likely RNAPIII-specific due to their higher similarity with promoters of the other RNAPIII snRNAs (U6, U6atac, etc.).

**Figure 5.**
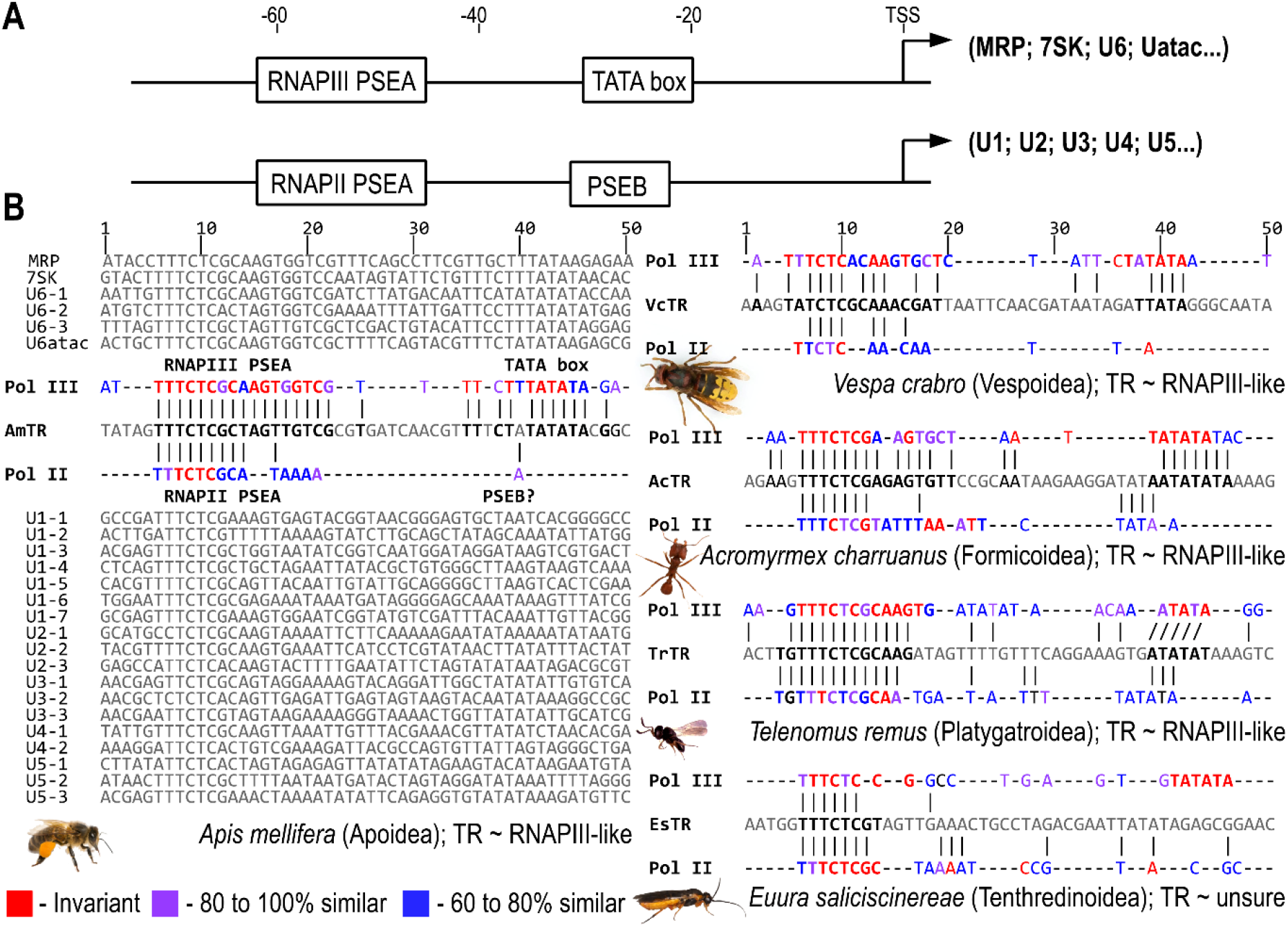
Comparison of the TR promoter with typical type 3 promoters of snRNAs. (A) Structural characteristics of the insect type 3 promoter for recognition by RNAPII or RNAPIII transcription factors from (35). (B) Multiple sequence alignments of type 3 snRNA promoters of snRNAs (from Table S3) and their comparison with TR promoter regions in five example species across Hymenoptera phylogeny. Except for the previously characterized type 3 promoter in *Apis mellifera* (35), presented as a full alignment of respective snRNA promoters, only consensus sequences (annotated as “Pol II” and “Pol IIl”) generated in the same manner for RNAPII resp. RNAPIII transcribed snRNAs are presented for other species.

### Structural features of Hymenopteran telomerase RNAs

We next determined if these putative TRs contained the structural domains known to be necessary for function in other eukaryotic lineages (36–42). TRs typically contain a core region comprised of a pseudoknot (PK) domain 3’ of the template, a template boundary element (TBE, also designated as P1.1) 5’ of the template, and a long range stem loop structure that closes this core region, known as the P1c domain. Using multiple sequence alignment guided structural predictions from the Meliponini (*n* = 10 representatives) and Bombus (n = 29) clades, which last shared a common ancestor ∼40 MYA, we identified the expected PK, TBE, and P1c domains (Figures 6A and B). The PK itself is typically comprised of long range interactions forming two stems (P2/P3) separated by loops of varying length. Within the Meliponini, the P2/3 stems are invariant, with the exception of a non-pairing nucleotide which is conserved in 90% of sampled taxa and retains its non-pairing state in the remaining taxa (Figure 6A). In the Bombus lineage, the P2/3 stems also display high conservation, with the exception of the non-pairing nucleotide within P3 that is positionally conserved between Bombus and Meliponini (Figure 6B). For both of these lineages, the P2 loop (J2/3_u_) is longer than the P3 loop, and contains the invariant U residues known to be essential for telomerase activity in vertebrates (Figures 6A and B). These essential motifs are present in each of the other sampled lineages (Figures 6C-F). Interestingly, outside of these essential motifs, other structural elements within the core region are not conserved between the sampled lineages, including multiple stem loops situated 3’ of the template and 5’ of the PK (Figure 6A-F). We observed strong evidence of conservation or covariation within the PK, TBE, and P1c in all sampled lineages with the exception of the Vespoidea (Figure 6D). Within the Vespoidea, 16 taxa were sampled from four genera (**Supplementary Figure S4**). While the P1c and P1.1 elements displayed evidence of strong conservation, the PK and P2.1 domains were each genera-specific, although positionally conserved within the primary sequence of the putative TRs, with a long P2.1 motif for all genera (Figure 6D, blue dashed boxes). Whether this reflects the evolutionary distance separating these genera, or if novel TRs were co-opted is unclear. Finally, the variation in TR length between sampled lineages meant that some of the putative TRs, particularly those in the Tenthredinoidea, consisted entirely of the core template + PK + P1c region (**Supplementary Figure S4**), with no evidence for P4-P6 domains observed in other eukaryotic lineages. In sum, our phylogenetically informed structural prediction of Hymenopteran TRs uncovered evidence for all of the domains necessary for telomere elongation, adding further support to these RNAs being *bona fide* TRs.

**Figure 6.**
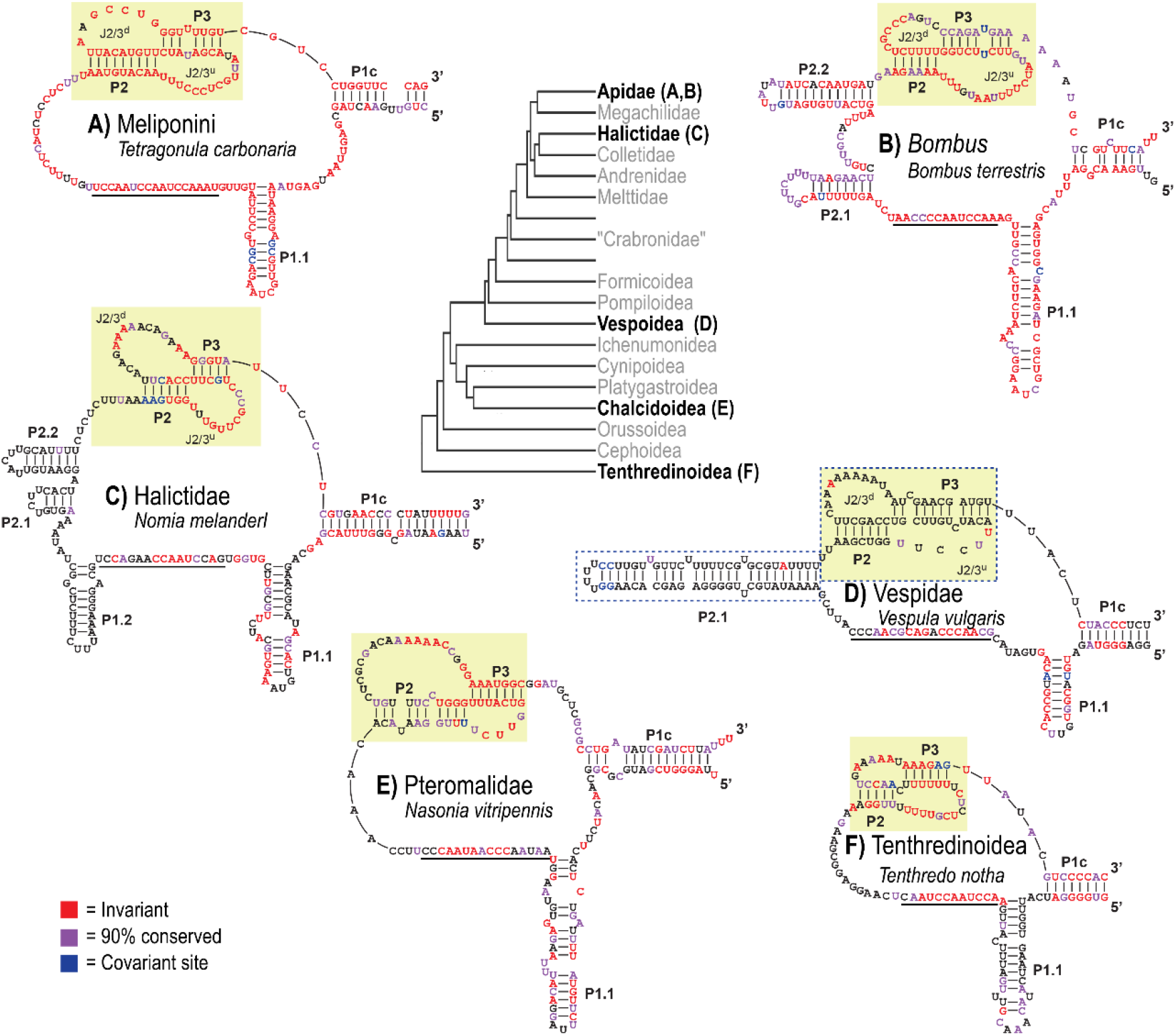
Comparative phylogenetic structural predictions across putative TRs in Hymenoptera uncover conserved essential core functional domains. (A) Predicted core Template/Pseudoknot (TPK) domain of Meliponini putative TRs (n = 10) using the *Tetragonula carbonaria* TR as reference. Template boundary element (TBE) is denoted as P1.1, the predicted template domain is underlined, the PK is denoted by the yellow box, and the stem loop closing the TPK is denoted as P1c. Invariant nucleotides are red, sites with >90% conservation (within the sampled lineages) are purple, and sites with evidence of covariation are blue. (B-F) Predicted core TPK domains for Bombus, Halictidae, Vespidae, Pteromalidae, and Tenthredinoidea, respectively. Species used as references are denoted. The blue dashed boxes in (D) denote analogous structural features that are present in to major genera within the Vespidae that, while not sequence conserved, are structurally conserved and hence denoted as large-scale co-variation. Multiple sequence alignments supporting these structural predictions are found in **Supplementary File S4**.

### Is the type 3 snRNA promoter evolutionary conserved in the Arthropoda TRs?

The surprising switch to a type 3 snRNA promoter, presumably recognized by RNAPIII in Hymenoptera TRs, represents an unprecedented change in the molecular nature of TRs within animals. This type of TR is characteristic among different eukaryotes from the phylogenetic megagroup of Diaphoretickes, but has not been reported within animals, where TRs, as RNAPII transcribed H/ACA box RNAs, seem to be ancestral (6). We hypothesize that some dramatic evolutionary change from the ancestral *animal-like* TR (6) is associated with the divergence of Superphyllum Ecdysozoa involving primarily Nematodes and Arthropods, Nevertheless, we lack sufficient TR-specific knowledge despite previous attempts (6). To prove such a hypothesis, characterisation of TR subunits in these clades is needed. Species with relatively conserved type 3 promoters (resp. PSE sequence) can be utilised for prediction of a limited number of TR-like loci based on the PSE consensus sequence and template-like region situated downstream of the PSE. Subsequently, evolutionary conservation of predicted TR-like loci across related genomes may support or filter out particular TR candidates in a manner similar to that previously predicted for TRs across the eukaryote megagroup Diaphoretickes (4). To predict a relevant number of such TR candidates we needed: (i) a small genome; (ii) highly conserved PSE; (iii) known telomere motif; (iv) available genomes from both close and distant relatives. Here we test whether the Type 3 promoter is conserved in TRs in other clades outside Hymenoptera. To infer PSE sequence consensus, we identified typical type 3 promoter snRNAs (U1;…;U6; MRP;7SK) across all Arthropoda representative genomes (raw snRNA sequences extended with promoters are referred in **Supplementary Table S3**). Subsequently, snRNA sequences (particularly their promoter regions) were then screened using the MEME tool (21) and/or aligned in Geneious software to check the presence of shared sequence motifs among snRNAs tested within species analysed. We succeeded primarily in certain genomes from Lepidoptera, in which several genomes showed highly conserved PSE, including a previously characterized type 3 promoter in *Bombyx mori* (22)) (see example in **Figure 7B**). In most of the Insecta genomes, it was possible to predict thousands of sequences starting with PSE and bearing an insect type template-like region up to 200nt downstream of the PSE. The difficulty was to support any of these loci as the putative TR. First, such TR candidates should have homologs in other related genomes in which the template region and type 3 promoter are conserved. In contrast to other Insect clades, Lepidoptera with its almost 900 available representative genomes met our criteria for TR prediction (i – iv; mentioned above), and provided a robust space to support TR candidates in homology searches. Here we present the sequence and structure of the type 3 promoter snRNA (see examples in **Figure 7** and **Supplemementary Table S4**) bearing a template-like region which was conserved across Lepidoptera and Trichoptera genomes (note: Trichoptera genomes showed a low conservation of the type 3 promoter). Unfortunately, Lepidoptera and Trichoptera lacked any variability in telomere motif sequence, thus we were not able support this TR candidate by comparison with corresponding mutations in TR template regions and mutated telomere motifs. This was in contrast to Hymenoptera TRs, where numerous changes in the TR template were confirmed by the changed telomere sequence. Therefore, in this study, it was premature to assess whether this snRNA, conserved across Lepidoptera and Trichopterea phylogeny, acts as a genuine TR, or has a different function. On the other hand, these putative TRs displayed all of the key core structural elements, including a TBE and deeply conserved PK (Figure 7C) supporting their functionality as a TR genes.

**Figure 7.**
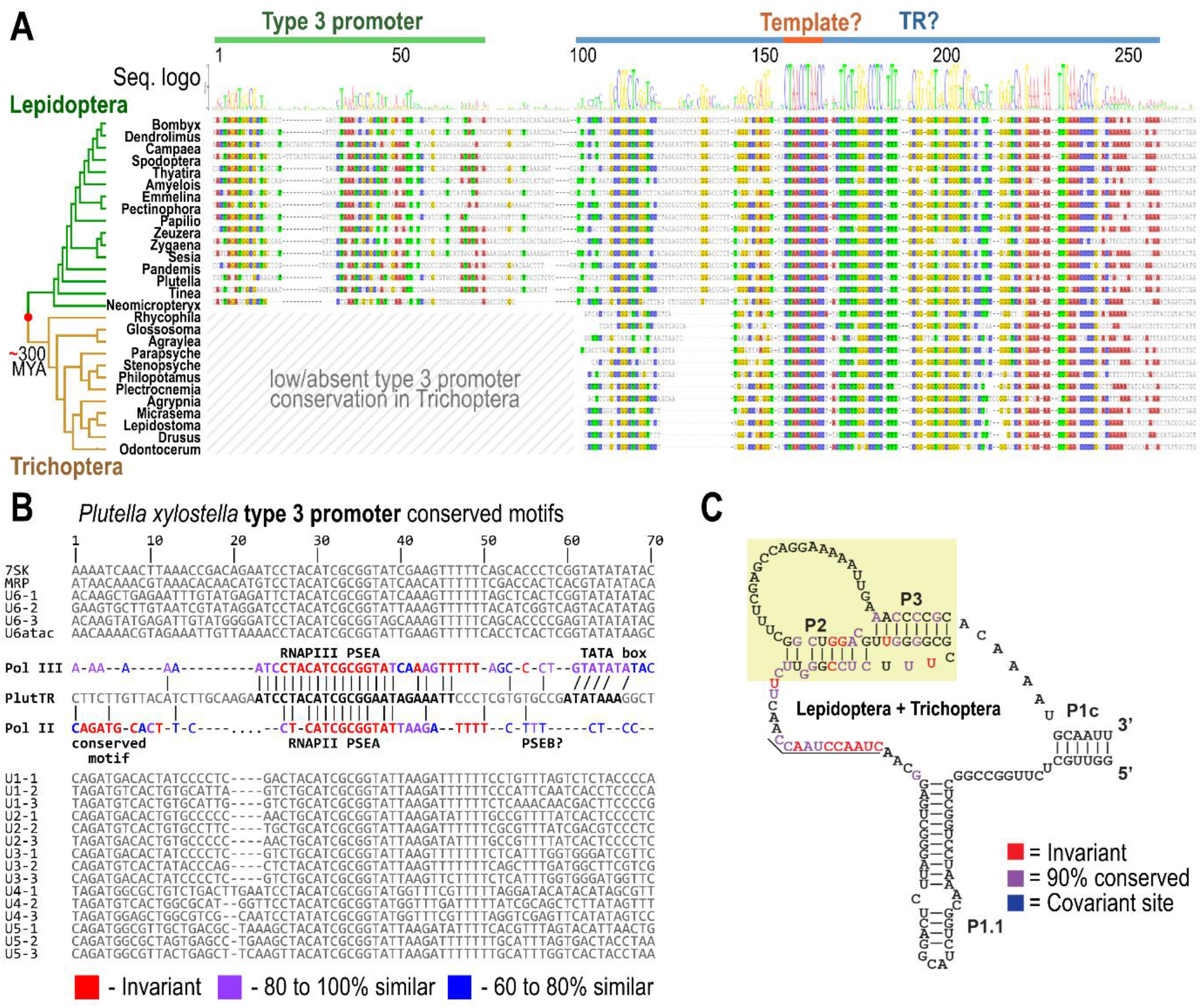
Evolutionary conservation of TR-like sequence homologs in Lepidoptera and Trichoptera harbouring type 3 promoter. (**A**) Sequence alignment of TR-like homologs in examples across Lepidoptera and Trichoptera phylogeny according to (43,44) (all TR-like homologs extended with promoter regions are referred in Supplementary Table S4). Conserved nucleotides (threshold ≥75%) are highlighted in colour. Promoters, TR-like sequences and their templates are annotated above the alignment. (**B**) Example of a comparison between promoter regions of the typical type 3 promoter snRNAs and the promoter of TR-like sequences in *Plutella xylostella* (top three snRNA sequences per snRNA species from **Supplementary Table S3** were included). Identified promoter motifs are highlighted in bold and coloured according to sequence similarity in consensus sequences corresponding to RNAPII or RNAPIII transcribed snRNAs (consensus sequences named as Pol II and Pol III respectively). If RNAPII or RNAPIII act on TR-like sequences in Lepidoptera remains unclear (C) Consensus structure of TR-like genes in Lepidoptera and Trichoptera suggesting presence and conservation of domains P1c; TBE (P1.1), Template, and PK (P2+P3) proposed according to a multiple sequence alignment (**Supplementary Figure S4**). Invariant sites are shown in red, sites with >90% conservation shown in purple and covariant sites are shown in blue.

## CONCLUSIONS

Our results reveal TRs in the most divergent animal phylum – Arthropoda, more precisely in the insect orders of Hymenoptera and Lepidoptera. Contrary to all other TRs described within the Animalia kingdom so far, the newly characterised TRs are snRNAs transcribed by RNAPIII under the control of the type 3 promoter – the type of TRs recently described in plants and other taxa across the phylogenetic megagroup Diaphoretickes (4). These results demonstrate a secondary evolutionary switch from the ancestral Animalia H/ACA box RNAPII transcribed snoRNAs, which was associated with the divergence of Arthropoda. Our findings provide an explanation of previous failures to identify TRs in insects. We further identify the Cynopoidea lineage in Hymenoptera, in which telomerase has probably been replaced with a telomerase-independent system of telomere maintenance – similarly to the earlier described cases in Drosophila and some other Diptera (1,28). The switch to TRs of the snRNA-type, or the switch to telomerase-independent telomere maintenance could thus represent examples of alternative evolutionary solutions of the loss of function of the ancestral telomerase system in some Animalia taxa.

## SUPPLEMENTARY DATA

Supplementary data are available at BioRxiv online

## DATA AVAILABILITY

All Hymenopteran TR genes identified and inferred in this study are available in **Supplementary Table S1**, sequences are available at GenBank under accessions: BK062014,…,BK062274. Putative TR-like candidates in Lepidoptera and Trichoptera are referred to in **Supplementary Table S4**.

## ACKNOWLEDGEMENT

Computational resources were supplied by the project “e-Infrastruktura CZ” (e-INFRA CZ LM2018140) supported by the Ministry of Education, Youth and Sports of the Czech Republic. Computational resources were provided by the ELIXIR-CZ project (LM2018131), part of the international ELIXIR infrastructure. We acknowledge the CF Genomics supported by the NCMG research infrastructure (LM2018132 funded by MEYS CR) for their support with obtaining scientific data presented in this paper. We would like to thank Gary Blissard (BTI, Cornell Univ.) for helpful discussions.

## REFERENCES

1. Biessmann, H. and Mason, J.M. (2003) Telomerase-independent mechanisms of telomere elongation. Cellular and Molecular Life Sciences, 60, 2325–2333.

2. Podlevsky, J.D. and Chen, J.J.L. (2016) Evolutionary perspectives of telomerase RNA structure and function. Rna Biology, 13, 720–732.

3. Fajkus, P., Peska, V., Zavodnik, M., Fojtova, M., Fulneckova, J., Dobias, S., Kilar, A., Dvorackova, M., Zachova, D., Necasova, I. et al. (2019) Telomerase RNAs in land plants. Nucleic Acids Research, 47, 9842–9856.

4. Fajkus, P., Kilar, A., Nelson, A.D.L., Hola, M., Peska, V., Goffova, I., Fojtova, M., Zachova, D., Fulneckova, J. and Fajkus, J. (2021) Evolution of plant telomerase RNAs: farther to the past, deeper to the roots. Nucleic Acids Research, 49, 7680–7694.

5. Waldl, M., Thiel, B.C., Ochsenreiter, R., Holzenleiter, A., Oliveira, J.V.D., Walter, M.E.M.T., Wolfinger, M.T. and Stadler, P.F. (2018) TERribly Difficult: Searching for Telomerase RNAs in Saccharomycetes. Genes, 9.

6. Logeswaran, D., Li, Y., Podlevsky, J.D. and Chen, J.J.L. (2021) Monophyletic Origin and Divergent Evolution of Animal Telomerase RNA. Molecular Biology and Evolution, 38, 215–228.

7. Shakirov, E.V., Chen, J.J. and Shippen, D.E. (2022) Plant telomere biology: The green solution to the end-replication problem. Plant Cell.

8. Greider, C.W. and Blackburn, E.H. (1989) A Telomeric Sequence in the Rna of Tetrahymena Telomerase Required for Telomere Repeat Synthesis. Nature, 337, 331–337.

9. Hargrove, B.W., Bhattacharyya, A., Domitrovich, A.M., Kapler, G.M., Kirk, K., Shippen, D.E. and Kunkel, G.R. (1999) Identification of an essential proximal sequence element in the promoter of the telomerase RNA gene of Tetrahymena thermophila. Nucleic Acids Research, 27, 4269–4275.

10. Lingner, J., Hendrick, L.L. and Cech, T.R. (1994) Telomerase Rnas of Different Ciliates Have a Common Secondary Structure and a Permuted Template. Gene Dev, 8, 1984–1998.

11. Burki, F., Roger, A.J., Brown, M.W. and Simpson, A.G.B. (2020) The New Tree of Eukaryotes. Trends in Ecology & Evolution, 35, 43–55.

12. Lukhtanov, V.A. (2022) Diversity and evolution of telomere and subtelomere DNA sequences in insects. bioRxiv, 2022.2004.2008.487650.

13. Zhou, Y.H., Wang, Y., Xiong, X., Appel, A.G., Zhang, C. and Wang, X. (2022) Profiles of telomeric repeats in Insecta reveal diverse forms of telomeric motifs in Hymenopterans. Life Science Alliance, 5.

14. Benson, G. (1999) Tandem repeats finder: a program to analyze DNA sequences. Nucleic Acids Research, 27, 573–580.

15. Peska, V., Sitova, Z., Fajkus, P. and Fajkus, J. (2017) BAL31-NGS approach for identification of telomeres de novo in large genomes. Methods, 114, 16–27.

16. Novak, P., Neumann, P. and Macas, J. (2020) Global analysis of repetitive DNA from unassembled sequence reads using RepeatExplorer2. Nat Protoc, 15, 3745–3776.

17. Nawrocki, E.P. and Eddy, S.R. (2013) Infernal 1.1: 100-fold faster RNA homology searches. Bioinformatics, 29, 2933–2935.

18. Peri, S., Roberts, S., Kreko, I.R., McHan, L.B., Naron, A., Ram, A., Murphy, R.L., Lyons, E., Gregory, B.D., Devisetty, U.K. et al. (2020) Read Mapping and Transcript Assembly: A Scalable and High-Throughput Workflow for the Processing and Analysis of Ribonucleic Acid Sequencing Data. Frontiers in Genetics, 10.

19. Raden, M., Ali, S.M., Alkhnbashi, O.S., Busch, A., Costa, F., Davis, J.A., Eggenhofer, F., Gelhausen, R., Georg, J., Heyne, S. et al. (2018) Freiburg RNA tools: a central online resource for RNA-focused research and teaching. Nucleic Acids Research, 46, W25–W29.

20. Griffiths-Jones, S., Bateman, A., Marshall, M., Khanna, A. and Eddy, S.R. (2003) Rfam: an RNA family database. Nucleic Acids Research, 31, 439–441.

21. Bailey, T.L., Johnson, J., Grant, C.E. and Noble, W.S. (2015) The MEME Suite. Nucleic Acids Research, 43, W39–W49.

22. Hernandez, G., Valafar, F. and Stumph, W.E. (2007) Insect small nuclear RNA gene promoters evolve rapidly yet retain conserved features involved in determining promoter activity and RNA polymerase specificity. Nucleic Acids Research, 35, 21–34.

23. Mei, Y., Jing, D., Tang, S.Y., Chen, X., Chen, H., Duanmu, H.N., Cong, Y.Y., Chen, M.Y., Ye, X.H., Zhou, H. et al. (2022) InsectBase 2.0: a comprehensive gene resource for insects. Nucleic Acids Research, 50, D1040–D1045.

24. Wickham, H. (2009) ggplot2 Elegant Graphics for Data Analysis Introduction. Ggplot2: Elegant Graphics for Data Analysis, 1-+.

25. Korandova, M., Krucek, T., Vrbova, K. and Frydrychova, R.C. (2014) Distribution of TTAGG-specific telomerase activity in insects. Chromosome Res, 22, 495–503.

26. Frydrychova, R. and Marec, F. (2002) Repeated losses of TTAGG telomere repeats in evolution of beetles (Coleoptera). Genetica, 115, 179–187.

27. Fajkus, P., Peska, V., Sitova, Z., Fulneckova, J., Dvorackova, M., Gogela, R., Sykorova, E., Hapala, J. and Fajkus, J. (2016) Allium telomeres unmasked: the unusual telomeric sequence (CTCGGTTATGGG)(n) is synthesized by telomerase. Plant Journal, 85, 337–347.

28. Mason, J.M., Randall, T.A. and Frydrychova, R.C. (2016) Telomerase lost? Chromosoma, 125, 65–73.

29. Prusakova, D., Peska, V., Pekar, S., Bubenik, M., Cizek, L., Bezdek, A. and Frydrychova, R.C. (2021) Telomeric DNA sequences in beetle taxa vary with species richness. Scientific Reports, 11.

30. Cervenak, F., Sepsiova, R., Nosek, J. and Tomaska, L. (2021) Step-by-Step Evolution of Telomeres: Lessons from Yeasts. Genome Biology and Evolution, 13.

31. Peska, V. and Garcia, S. (2020) Origin, Diversity, and Evolution of Telomere Sequences in Plants. Frontiers in Plant Science, 11.

32. Peska, V., Fajkus, P., Bubenik, M., Brazda, V., Bohalova, N., Dvoracek, V., Fajkus, J. and Garcia, S. (2021) Extraordinary diversity of telomeres, telomerase RNAs and their template regions in Saccharomycetaceae. Scientific Reports, 11.

33. Peska, V., Matl, M., Mandakova, T., Vitales, D., Fajkus, P., Fajkus, J. and Garcia, S. (2020) Human-like telomeres in Zostera marina reveal a mode of transition from the plant to the human telomeric sequences. Journal of Experimental Botany, 71, 5786–5793.

34. Koubova, J., Jehlik, T., Kodrik, D., Sabova, M., Sima, P., Sehadova, H., Zavodska, R. and Frydrychova, R.C. (2019) Telomerase activity is upregulated in the fat bodies of pre-diapause bumblebee queens (Bombus terrestris). Insect Biochemistry and Molecular Biology, 115.

35. Hernandez, N. (2001) Small nuclear RNA genes: a model system to study fundamental mechanisms of transcription. J Biol Chem, 276, 26733–26736.

36. Tzfati, Y., Fulton, T.B., Roy, J. and Blackburn, E.H. (2000) Template boundary in a yeast telomerase specified by RNA structure. Science, 288, 863–867.

37. Chen, J.L. and Greider, C.W. (2003) Template boundary definition in mammalian telomerase. Gene Dev, 17, 2747–2752.

38. Tzfati, Y., Knight, Z., Roy, R. and Blackburn, E.H. (2003) A novel pseudoknot element is essential for the action of a yeast telomerase. Gene Dev, 17, 1779–1788.

39. Chen, J.L. and Greider, C.W. (2005) Functional analysis of the pseudoknot structure in human telomerase RNA. P Natl Acad Sci USA, 102, 8080–8085.

40. Cash, D.D., Cohen-Zontag, O., Kim, N.K., Shefer, K., Brown, Y., Ulyanov, N.B., Tzfati, Y. and Feigon, J. (2013) Pyrimidine motif triple helix in the Kluyveromyces lactis telomerase RNA pseudoknot is essential for function in vivo. P Natl Acad Sci USA, 110, 10970–10975.

41. Jansson, L.I., Akiyama, B.M., Ooms, A., Lu, C., Rubin, S.M. and Stone, M.D. (2015) Structural basis of template-boundary definition in Tetrahymena telomerase. Nat Struct Mol Biol, 22, 883–888.

42. Song, J.R., Logeswaran, D., Castillo-Gonzalez, C., Li, Y., Bose, S., Aklilu, B.B., Ma, Z.Y., Polkhovskiy, A., Chen, J.J.L. and Shippen, D.E. (2019) The conserved structure of plant telomerase RNA provides the missing link for an evolutionary pathway from ciliates to humans. P Natl Acad Sci USA, 116, 24542–24550.

43. Malm, T., Johanson, K.A. and Wahlberg, N. (2013) The evolutionary history of Trichoptera (Insecta): A case of successful adaptation to life in freshwater. Systematic Entomology, 38, 459–473.

44. Kawahara, A.Y., Plotkin, D., Espeland, M., Meusemann, K., Toussaint, E.F.A., Donath, A., Gimnich, F., Frandsen, P.B., Zwick, A., dos Reis, M. et al. (2019) Phylogenomics reveals the evolutionary timing and pattern of butterflies and moths. P Natl Acad Sci USA, 116, 22657–22663.

