## Supplementary Figures for "Hymenoptera (Insecta) telomerase RNAs switched to plant/ciliate-like biogenesis"

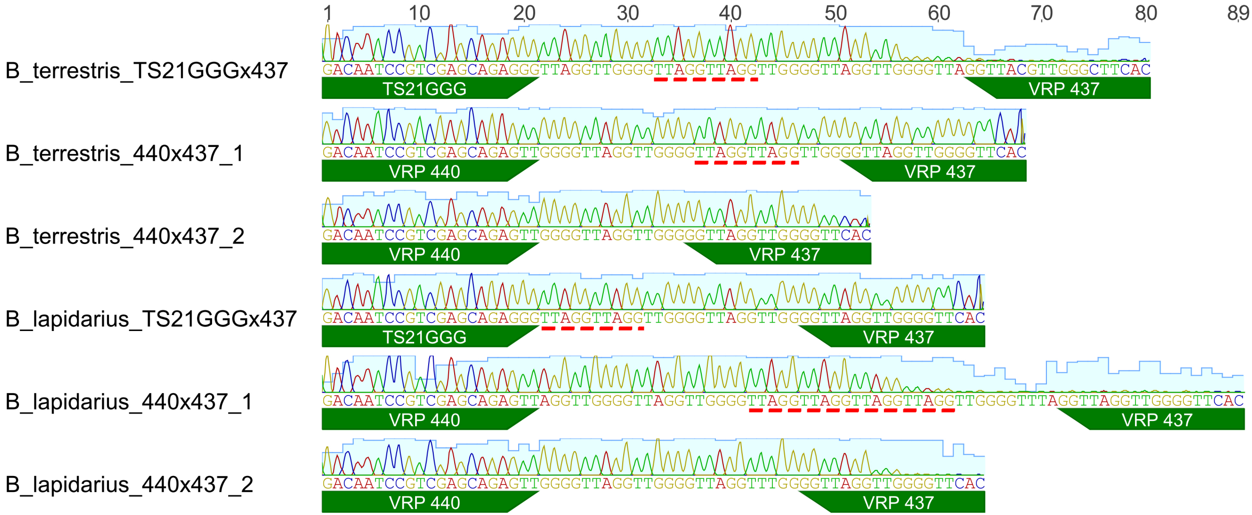


```
3'-CTAACCCCAATCCAAAG-5'
     ||---------->
5'-..ttGGGGTTAGGTT-3'

   CTAACCCCAATCCAAAG
         ||||---->
   ......ggttAGGTT

   CTAACCCCAATCCAAAG
        |||------>
   .....gggTTAGGTT
```

Analysis of the cloned TRAP products indicates variable **annealing** of *Bombus* **TR template** to substrate.

Besides dominant GGGGTTAGGTT telomere repeat in *Bombus*, $[\text{TTAGG}]_{2\text{-}3}$ was frequently present in **extension products**.

**Suppementary Figure S1:** Cloned PCR products from TRAP assay in *B. terrestris* and *B. lapidarius* showed synthesis of TTAGGTTGGGG repeat motif by telomerase. Frequently observed [TTAGG]2-3 repeats (red underlined) presumably correspond to alternative annealing of TR template to substarte primer or extension product resulting in synthesis of an incomplete telomere repeat *in vitro*.

**Suppementary Figure S2:** TERT search in Hymenoptera.(A) Example alignment of TERT conserved motifs in representatives of Hymenoprea phylogeny. TERT full-length proteins were extracted from InsectBase 2.0 (Mei, 2022). (B) Search for TERT-like sequences in available representative Hymenptera genomes (at NCBI). Full length TERT protein sequences were split in to two parts (containing TRBD and RT domains respectively) and used as a queries (in row) in tbastn searches against representative genomes (in column). Blast hit significance (e-value) is distinguished in colour. In contrast to other clades, no TERT-like sequences were observed in Cynipoidea (Note: hits observed in tblastn with RT domain in Callyrthis sp. seem to be retroelement and not a single- or low- copy gene.)

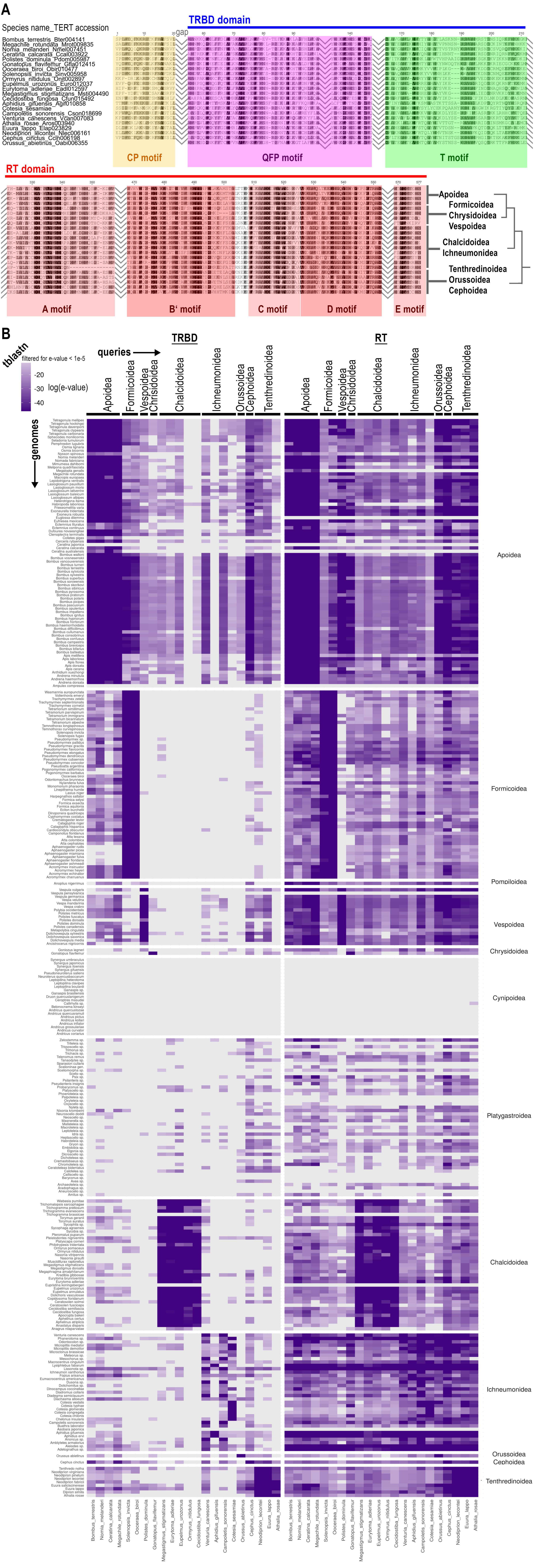

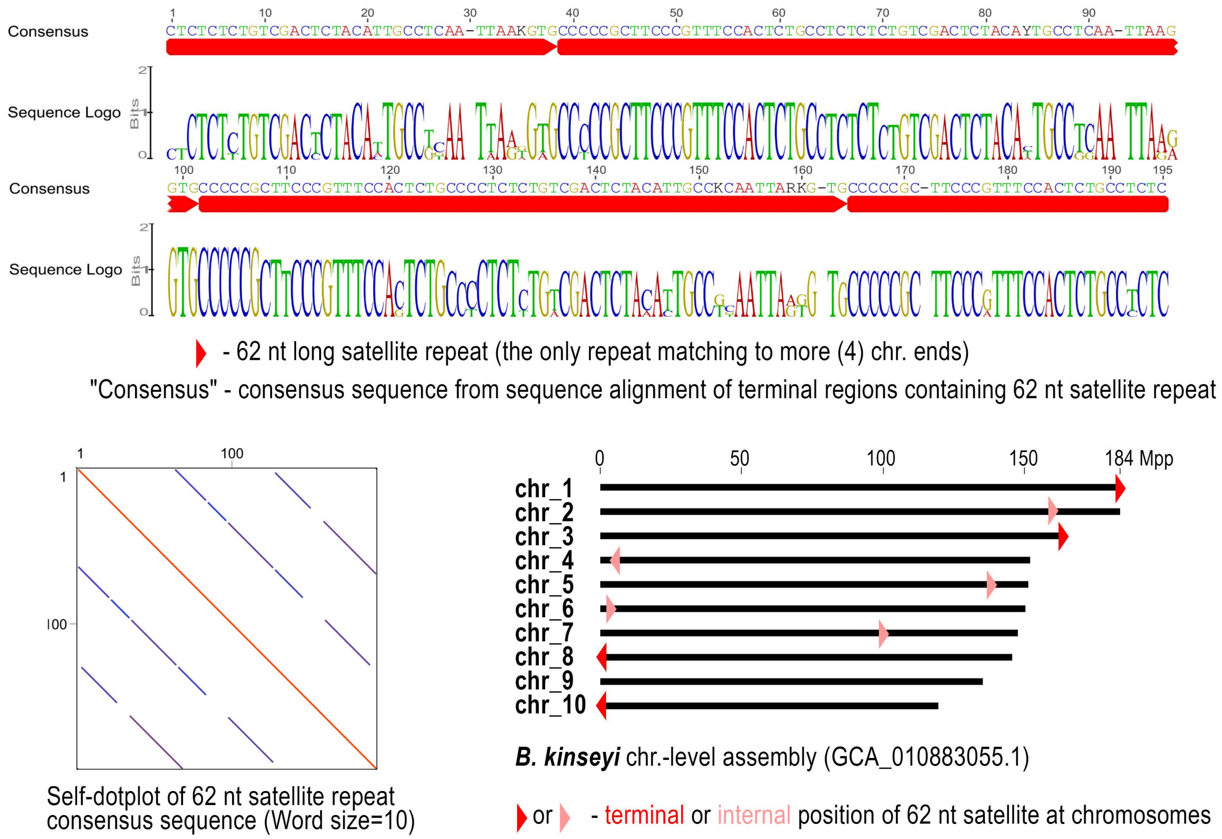


**Suppementary Figure S3:** Identification of common DNA sequences between *B. kinseyi* chromosome termini. 2kb long chromosome terminal sequences were compeared to each other by blastn.

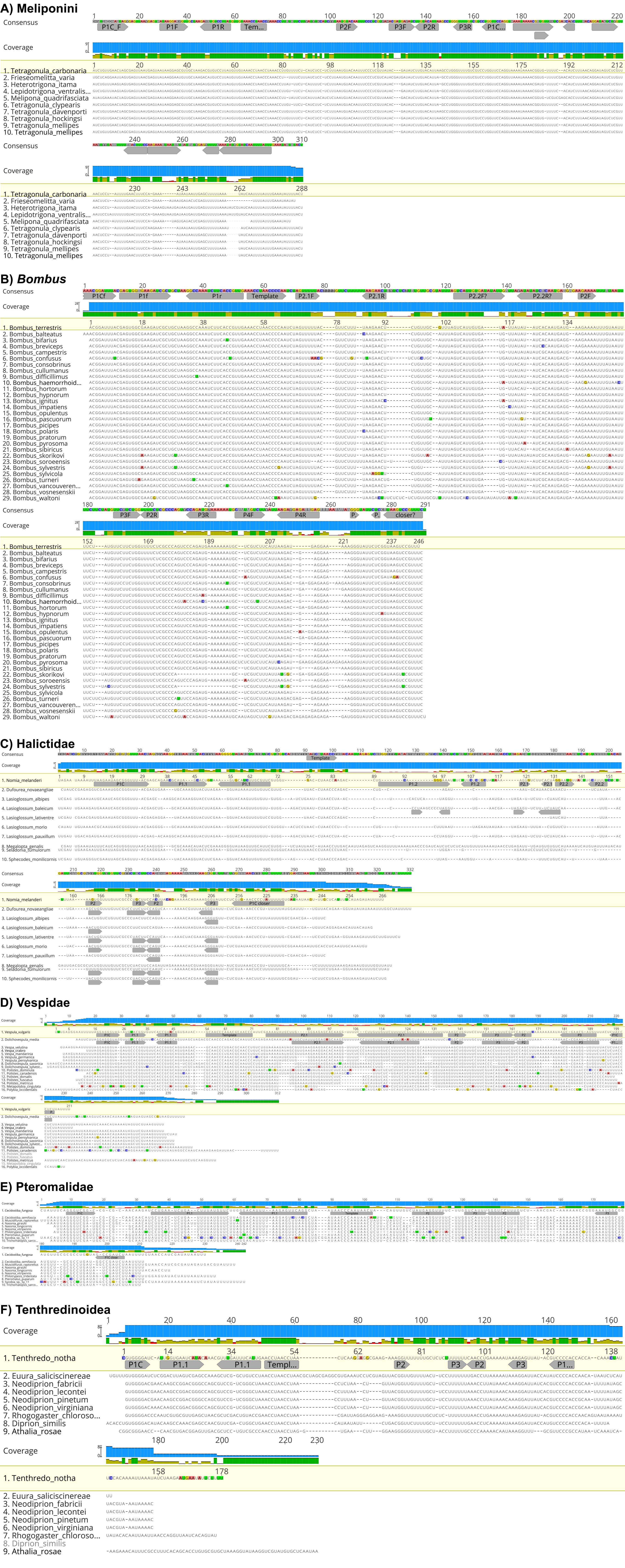


**Supplementary Figure S4:** Multiple sequence alignments (MSA) of TR genes from representatives of particular Hymenoptera clades (A-F). Conserved RNA secondary structre elements are depicted by grey arrows indicating mutualy paried RNA sequences. TR core structures inluding imprtant functional domains (for example, TBE, Template, PK) are shownin Figure 6.
